# Configural behavior response to a two component odor mixture

**DOI:** 10.1101/2023.08.20.552323

**Authors:** Rory F. O’Brien, Dmytro Klymyshyn, Lawrence B. Cohen

**Affiliations:** Department of Cellular and Molecular Physiology, Yale University School of Medicine, New Haven CT, USA

## Abstract

Configural odors are mixtures of odorant molecules that are perceived as an odor distinct from the components. They provide an attractive target for investigations of odor perception in the olfactory bulb. We investigated the perception of octanal and citronellal in mice using a Go/No-Go behavioral paradigm, and show that mice trained on the mixture of the two odorants respond selectively to the mixture, but not to the components delivered separately.

## Introduction

Configural odor perception refers to the perception of a mixture of odors as a separate odor, distinct from any of the component odors. This is opposed to elemental perception, in which individual component odors can be recognized. Most odor mixtures made up of more than a few component odorants are perceived as configural (Laing and Francis, 1989; Weiss et al, 2012), and indeed, many odors we encounter (such as flower smells) are composed of tens to hundreds of distinct odorant molecules (Howard & Gottfried, 2014; Lain & Francis, 1989) which form one configural percept.

Based on its ubiquity within our sensory experience, the phenomenon of configural odor perception is evidently important for olfactory comprehension of the world, and yet its implementation within the olfactory circuitry of the brain is not understood. Given the olfactory bulb’s intrinsic discrimination between odorant molecules at the level of glomeruli, which likely contributes to various features of olfactory perception such as adaptation and concentration invariance of odor recognition (e.g. Storace & Cohen, 2017, 2021), it is possible that configural perception could be an additional function of the olfactory bulb itself, perhaps together with feedback from higher olfactory processing centers.

Another possible explanation for configural odor perception is the receptor-level interactions among odorant molecules themselves. Each olfactory sensory neuron expresses one particular odor receptor, but each receptor responds to more than one molecule, with some being agonists and others antagonists, allosteric modulators, etc. (Xu *et al*., 2020) It is possible that this complex web of interactions between odorant molecules (and even non-odorant volatiles) across multiple receptors is responsible for more complex olfactory phenomena such as configural perception.

Determining the contribution of the olfactory bulb in generating configural perceptions will be simpler if the odorant mixture involves only two or three individual components. Table 1 lists five examples of two or three odorants that, when mixed in the correct concentration ratios, are reported to yield configural perceptions. Disconcertingly, an attempt to replicate the pineapple configural response using a different kind of olfactometer was not successful (Acree et al, 2014).

**Table 1.**

| Configural perception | Individual odorants | Animal | Reference |
| --- | --- | --- | --- |
| 1. unknown | octanal, citronellal | rat | Kay et al, 2003 |
| 2. unknown | eucalyptol, benzaldehyde | rat | Kay et al, 2005 |
| 3. potato chips | methanethiol, methional,<br>2-ethyl-3,5-dimethylpyrazine | human | Rochelle et al, 2018 |
| 4. butter | diacetyl, butanoic acid,<br>decalactone | human | Schieberle et al, 1993 |
| 5. pineapple | ethyl isobutyrate, ethyl maltol<br>allyl-α-ionone | human<br>rabbit | Le Berre et al, 2008<br>Coureaud et al, 2008 |

We have carried out behavioral experiments on mixture 1 using concentration ratios shown to give configural or component responses (Kay et al, 2003). Although the odorant concentration ratios for generating the rat’s configural response are reported, the actual odorant concentrations were not determined.

Mice were used in this study due to future plans to incorporate optical measurement of activity in different transgenically-labeled cell types in the olfactory bulb of head-fixed animals. For other species which have previously been used in olfaction experiments, the genetic tools necessary to carry out this sort of experiment do not exist, so we have used mice to establish a baseline for future research.

We used a Go/No-Go task to train discrimination between this configural mixture and a third. We investigated the configurality of the odor mixture by presenting the component odorants to animals trained on different ratios. We report a configural response to a 1:1 ratio of citronellal to octanal, but an elemental response to a 30:1 ratio.

## Methods

All experiments were performed in accordance with relevant guidelines and regulations, including a protocol approved by the Institutional Animal Care and Use Committee at Yale University. Mice were housed under standard environmental conditions under a 12h light/dark cycle. Our measurements were made during the light phase.

During behavioral testing, the mice were head-fixed and their bodies held within a plastic tube. A Sanworks Bpod 2.0 state machine was used to detect licks and coordinate rewards. Following Komiyama et al., (2010) and Guo et al., (2014), we trained head fixed mice in a Go/No-Go task to lick in response to a mixture of citronellal and octanal (Go) and not lick in response to hexanol (No-Go). The mice were water-deprived and completed 150-300 trials each day.

The trial timeline is shown in Figure 1. An LED turned on at trial start to prime the mouse for the odor delivery. Then, after a delay of 1 sec from the odor delivery, water would either be delivered (on Go trials) or not delivered (on No-Go trials). The mouse then had a fixed decision period of 1 sec in which to either lick or not lick. A correct response (lick for citronellal/octanal or no lick for hexanol) would end the trial after a 3 second delay, whereas an incorrect response would impose an additional 5 second time-out penalty period.

**Figure 1.**
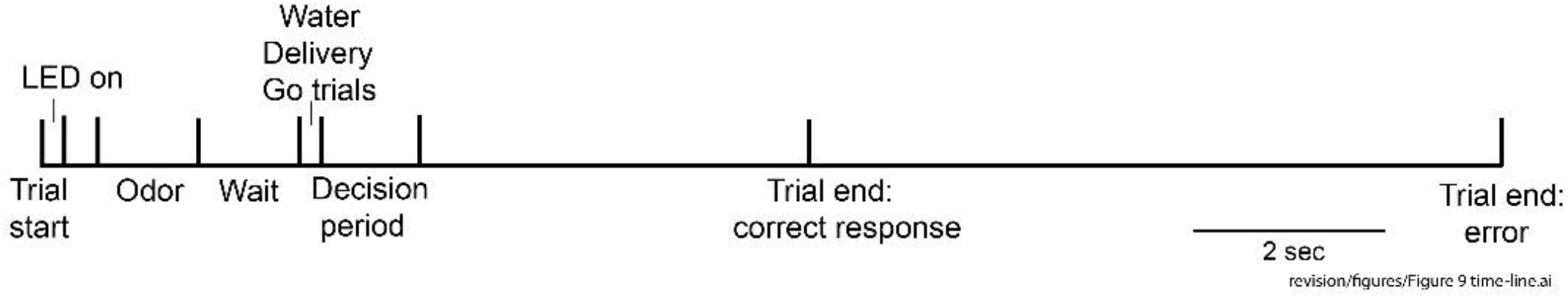
Go/No-Go behavioral trial design.

Interspersed with training trials were test trials. In these trials, the Go mixture was presented, but no water. If the mice licked when the Go odorant mixture was presented without water, we concluded that they had learned to lick in response to the Go mixture, and were not simply detecting the water delivery.

The olfactometer followed that described in Vučinić *et al*. (2006), such that introducing odorant vapor into the air flow did not alter the air flow; it displaced an equal volume of diluent air. This ensured that the rate, humidity, and direction of airflow remained constant to avoid confounding the olfactory stimulus with a purely mechanical stimulus derived from changes in the properties of the air flow.

## Results

As illustrated in Figure 2A and 2B, mice quickly learned to lick for the Go odor mixture, while learning to not lick for the No-Go mixture took longer. They begin with a high proportion of licks on Go trials, but also a relatively high (albeit substantially lower) proportion of licks on No-Go trials, only developing proper discrimination (a low proportion of false alarms) later in the first training day and on later training days.

**Figure 2.**
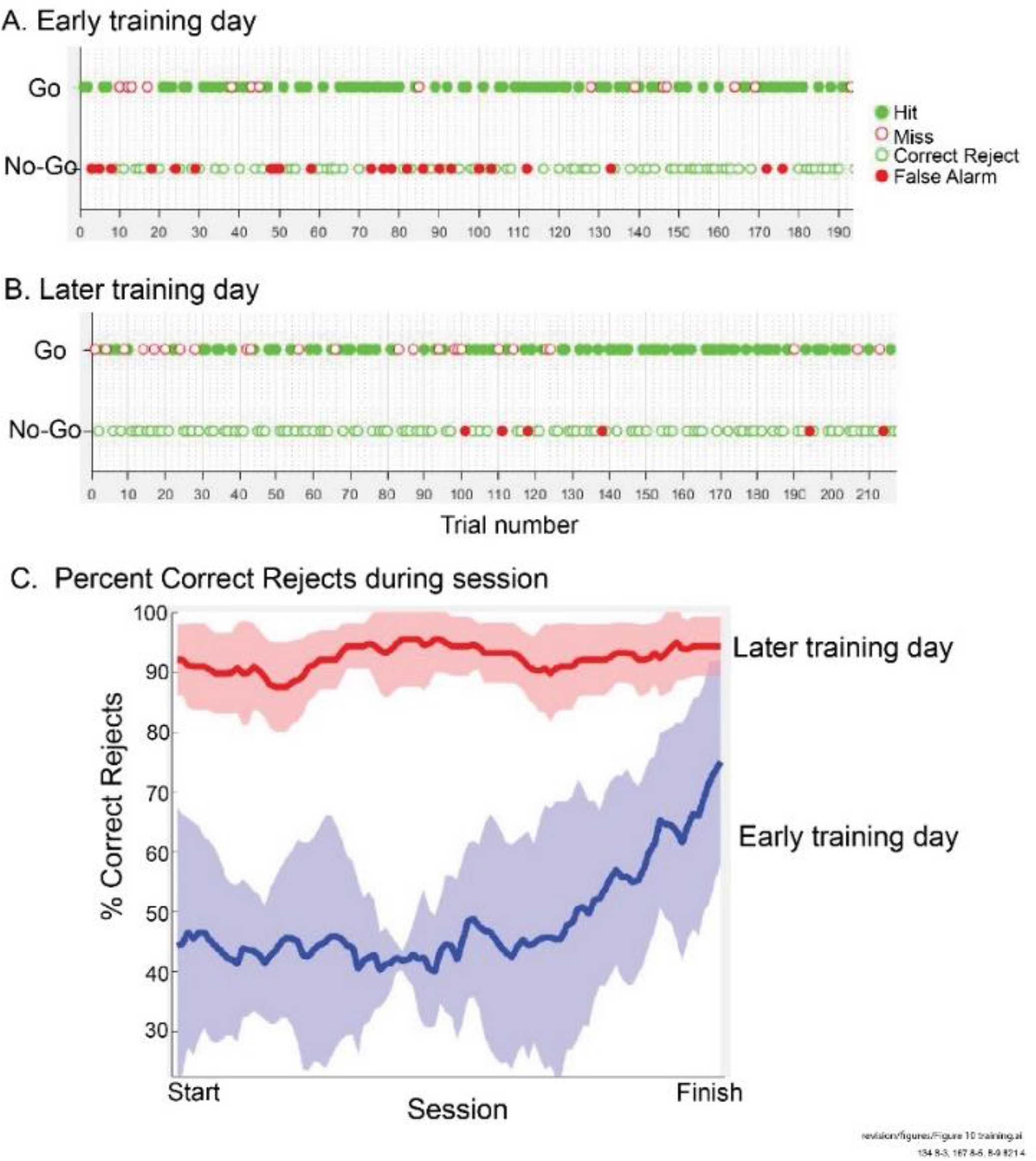
A: Performance of a representative mouse on an early training day, indicating a high proportion of False Alarm responses early in the session. B: Performance of the same mouse on a later training day, demonstrating learning of the Correct Reject response. C: Correct Reject percentages over the course of training on Early and Later training days (21-trial rolling average, shading represents 2 times SEM).

Because mice learned the correct Go response quickly, following Komiyama et al., (2010), we used Correct Rejects as a measure of learning. On an early training day the average of four mice achieved a 70% Correct Reject response by the end of their trials (Figure 2C, top). On a later training day they were ∼90% correct during the entire session (Figure 2C, bottom), results that were similar to those reported by Komiyama et al., (2010).

The response to the Go odor when the mouse is trained on a 1:1 mixture is configural, as shown in Figure 3A. The mouse does not respond to either of the component odorants, but it does respond to the combined odor, whether or not it is presented alongside water. This indicates that the odor of the test mixture produces a different olfactory percept than that produced by either of its two components – that is, it is configural.

**Figure 3.**
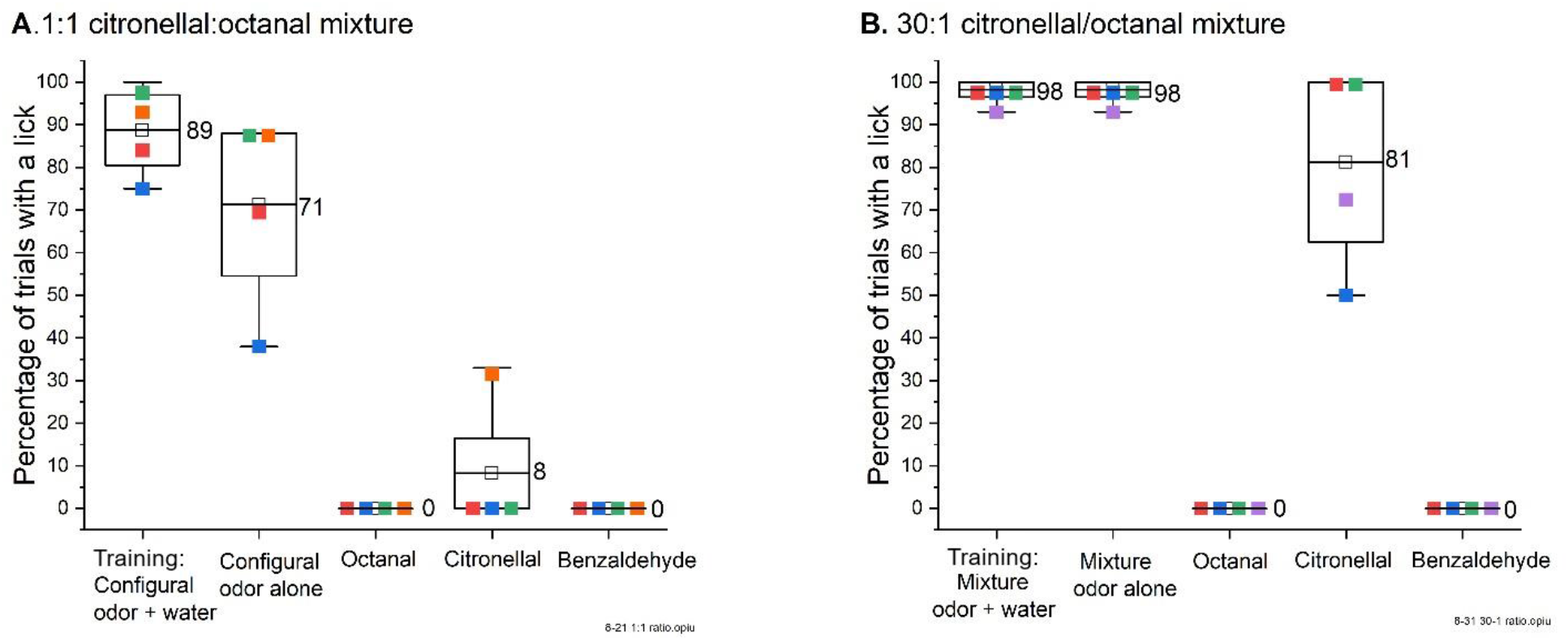
A: Comparison of licking percentage for different odors in mice trained on a 1:1 mixture of citronellal and octanal that was predicted to be configural. B: Comparison of licking percentage among different odors for mice trained on a 30:1 mixture of citronellal and octanal that was predicted to be elemental.

This configural response was not obtained when the mixture was instead 30:1 citronellal/octanal (Figure 3B), as the mice trained on this mixture selectively responded to the stronger of the two elemental odors (in this case, citronellal). This confirms the results obtained with the same odorant ratios in free moving rats in Kay *et al*., 2003.

As shown in Figure 3, the mice responded as well to the Go mixture alone as to the Go mixture with water. This indicates that they respond primarily to the odor cue, and not to the delivery of water itself.

## Discussion

The presented behavioral paradigm provides a robust platform on which the role of the olfactory bulb in configural odor perception can be studied using an imaging-compatible behavioral task which can be easily repeated hundreds of times in a session. Our data show that mice learn this task quickly, reaching substantial proficiency by the end of their first training session. They also maintain nearly perfect accuracy over the course of later sessions, confirming the results obtained in a previous study using the same odorants (Kay *et al*., 2003).

Further, we verified that most mice do not “cheat” on this task by detecting the presence of water and licking for that rather than detecting the Go odor mixture. We also show that that their perception of the 1:1 citronellal:octanal mixture is indeed configural.

